# Stable changing fluid shear stress promotes osteogenesis

**DOI:** 10.1101/402651

**Authors:** Linyuan Wang, Fang Xiao, Lei Yu, Hang Su, Yueqin Yang, Song Wang

## Abstract

**Objective:** Bone marrow cells encounter various chemical and mechanical stimuli from the internal environment. In vivo, fluid shear stress (FSS) is one of the primary mechanical stimuli that affect bone marrow-derived mesenchymal stem cells (BMCs) activity. Since various cases of FSS influence BMCs activity differently, the purpose of this study is to determine how BMC activity in osteogenesis and osteoclastgenesis is affected by stable and unstable changing FSS.

**Method:** BMCs samples from the femur of the mouse, divided in to three groups: stimulate by stable changing FSS, unstable changing FSS, and no FSS. RT-PCR would be applied to detect OPG, RANKL, ALP, OCN, RUNX2, and RANK of all the samples at the 3^rd^ day. Alizarin red staining and TRAP staining would be applied to test all the samples at the 6^th^ day.

**Results:** The S group samples showed the lowest level in RANKL mRNA and showed the highest level in OPG mRNA. RANKL/OPG mRNA in the S group was the smallest during three groups. Comparing with the C group samples, RUNX2 mRNA in the S group was increased significantly. RANK mRNA in the C group was three times more than S group. S group had the largest area of mineralized nodules, and largest number of area ≥100µm, ≥500µm^2^ or ≥1000µm^2^. The area of positive TRAP stain in S group was the smallest among three groups.

**Conclusion:** Stable changing FSS significantly increases osteogenesis relating to OPG-RANKL-RANK pathway. Compared with unstable changing FSS, stable changing FSS afford a more appropriate stimulation to osteogenesis.

**Abbreviations:**

ALP: Alkaline phosphatase
BMCs: Bone marrow-derived mesenchymal stem cells
Cx43: Connexin 43
C3a: Complement Component 3a
JNK: c-Jun N-terminal kinase
ERK: Extracellular regulated protein kinases
ERK5: Extracellular-signal-regulated kinase
FSS: Fluid shear stress
FasL: First apoptosis signal Ligand
GSK3β: Glycogen synthase kinase 3β
LPA: Lysophosphatidic acid
MSCs: Mesenchymal stem cells
M-CSF: Macrophage colony-stimulating factor
MCP-1: Monocyte chemoattractant protein-1
RANK: Receptor Activator of Nuclear Factor κ B
RANKL: Receptor activator of nuclear factor κ B ligand
RUNX2: Runt-related transcription factor 2
P2Y_2_R: Receptor of P2Y_2_
LGR4: Repeat-containing G-protein-coupled receptor 4
Sema: Semaphorins
TRAP: Tartrate-Resistant Acid Ahosphatase
TRPM7: Transient receptor potential cation channel subfamily M member 7
TRAFs: Tumornecrosisfactorreceptor associatedfactors
OPG: Osteoprotegerin
OCN: Osteocalcin
OPN: Osteopontin

## 1. Introduction

Bone marrow-derived mesenchymal stem cells(BMCs) have been attracted considerable attention because of their multiple differentiation potential^[1]^. At bone tissue, osteoclasts and osteoblasts are the bone resorbing cells and bone forming cells respectively. They work with each other during the process of bone remodeling. Osteoclasts differentiate from mononuclear cells of the monocyte/macrophage lineage. Osteoblasts are derived from mesenchymal stem cells (MSC). Also, their differentiation and maturity are influenced by bone marrow environment. In vivo, bone cells encoutner different stimuli including chemical and mechanical ways^[2]^. The primary mechanical stimulation is fluid shear stress (FSS)^[3]^.

Research had found connection between FSS and osteogenesis or osteoclastgenesis. P_2_Y_2_R^[4]^, TRPM7^[5]^, Cx43 ^[6]^ and Megakaryocytes^[7]^ as mechanosensitive factors receive shear stress stimuli and then modulate osteogenesis. Osteoblast and osteoclast differentiation can be promoted from shear stress through the ERK5 pathway^[8–10]^. MLO-Y4 osteocyte-like cells were exposed to fluid flow shear stress in different period. OPG mRNA level was significantly upregulated at the first hour after FSS stimulation^[11]^. BMCs from 6-month-old Sprague Dawtey rats vertebrae were induced by 1,25(OH)D_3_ and intervened by FSS at 30min. TRAP activity was enhanced and bone resorption area was increased^[12]^. Chenglin Liu found that fluid flow induce calcium respond increasing in osteoclast ^[13]^.

Bone is a porous tissue. Mechanical loading lead to deformation, strain gradient and local pressure in bone tissue. It can drive the interstitial fluid flow and then causes FSS on bone marrow cells^[14]^. In vivo, FSS varies between 8-30 dyn/cm^2^ according to different physical activities^[15]^. Bone was compressed 2000 or 3000 micro-strain in vivo^[16]^. Verbruggen SW predicted that average interstitial fluid velocities is 60.5μ m/s and average maximum shear stresses was 11 Pa surrounding osteocytes in vivo^[3]^.

A suitable device to mimic the real internal environment in bone tissue is crucial for researching the specific mechanism between FSS and osteogenesis. In our studie, we used an oscillating like “see-saw” system that based on Xiaozhou Zhou `s mode^[17]^l to imitate FSS acting on BMCs in vivo. Compared with other FSS systems, this one was easier, cheaper and more convenient. There were two different FSS cases in our device. FSS can be divided into stable changing FSS and unstable changing FSS.

In this experiment, we supply two different FSS cases to BMCs and measured the marker factors of osteogenesis or osteoclastgenesis to analysis the influence on FSS to BMCs.

## 2. Materials and Methods

### 2.1 Cell Culture

Four weeks old male Kunming mice were sacrificed by breaking neck and the femurs were removed from the body. Specimens were taken off soft tissues and then rinsed in PBS. BMCs were harvested by injecting the marrow cavity of bones with complete culture medium (89% DMEM+10% fetal bovine serum+1% penicillin-streptomycin) after cutting off two ends of the bones. 2×10^6^ cells/well were cultured in the standard six-well plates with 2.5ml or 3.5ml complete culture medium. The complete culture medium was changed every 3 days.

### 2.2 Fluid Shear Stress device

The culture plate was a commercial six-well plate. The bottom area of each well was 9.6 cm^2^. And we set up 3 groups according to the different culture medium. In the C group (control group) there was 3.5ml culture medium in each well and keeped horizon. In the S group and V group, there were 3.5ml and 2.5ml medium in each well respectively. When the culture plate was located on the horizontal table concentrator, the depth of culture medium was about 2.604 mm and 3.6mm respectively.

The FSS protocol was produced by the devices in our lab. The motor device was a BETS-M5 table concentrator and shook at 0.5 Hz, 2 times a day for 30 minutes. The tray of the device was 240mm×230mm and the maximum inclination angle was 12°. The culture plates were located on the two sides of the tray and parallel to the roller of tray. When the table concentrator was inclining, the culture medium in every well was flowing from one side to the other side.

### 2.3 RNA Extractions and Transcript Quantification Using Real-Time PCR

After the BMCs were treated with FSS for 3 days, all of the group cells were washed with PBS twice. Total RNA was extracted by Trizol (Invitrogen, USA), Chloroform, isopropyl alcohol and ethanol in order, and then quantified (Kaiao, China) and reversed transcribed with RevertAid^TM^ First Strand cDNA Synthesis Kit (Fermentas, LTU). The cDNA was amplified with the incorporation of SYBR Green fluorescent nucleic acid stain (Invitrogen, USA) by Real-Time PCR machine (ABI, USA). 2^[-ΔΔcycle threshold (Ct)]^ value was used to calculate the fold change in the expression of cytokines. ΔCt=Ct of cytokines - Ct of â-actin; ΔΔCt=ΔCt of treated groups - ΔCt of control group. Primers used in the RT-PCR for each gene were shown in Table 1.

**Table 1.**
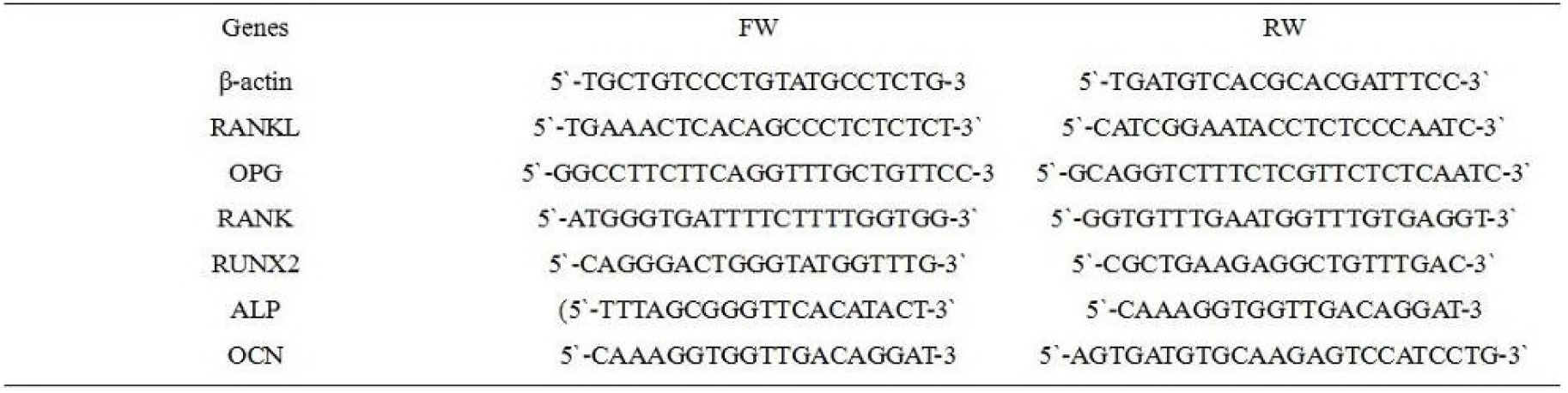
List of the primers used for the RT-PCR

### 2.4 Alizarin Red Staining

After the BMCs were cultured for 6 days, they were washed with the PBS twice. Ethanol was used to fix the cells and then 0.1% alizarin red S –Tris – Hcl (pH = 8.3) was used to stain the cells at 37°C for 30 minutes. Mineralized nodules were red in the microscope (Olympus, Japan). The number of mineralized nodules where the area is more than 100 μm^2^, 500 μm^2^ and 1000 μm^2^ of mineralize nodules and total area of it were calculated respectively (Image-Pro Plus 6.0).

### 2.5 TRAP Staining

After the BMCs were treated for 6 days, they were rinsed with PBS twice for 3 minutes. 2.5% glutaraldehyde was used to fix the cells and TRAP was used to stain the cells at 37°C for 50 minutes. The osteoclast cells were identified in the microscope as orange-red cells. Each sample was taken ten fields of view under microscope. The positive cell areas was calculated (Image-Pro Plus 6.0).

### 2.6 Statistics

The change of culture medium depth was evaluated using Pro-e 5.0 software. One-way ANOVA was used to assess differences among the groups by GraphPad Prism 5.0 software. The data was expressed as the mean ± standard deviation. Statistical significance was determined by *p<0.05*.

## 3. Results

### 3.1 Changing Fluid shear stress in three groups

The FSS pattern in a rocking dish was complex and spatially heterogeneous. For profiling the different conditions of the three groups, we set up a middle vertical cross-section of the culture medium in the well to quantify parameters. Cartesian coordinate system was created with X axis along dish bottom, Z axis along dish side wall, and the origin point (O) was located at the rotation center. The entire dish rotated in the vertical plane along pivotal point (O).(Figure 1)

**Figure 1.**
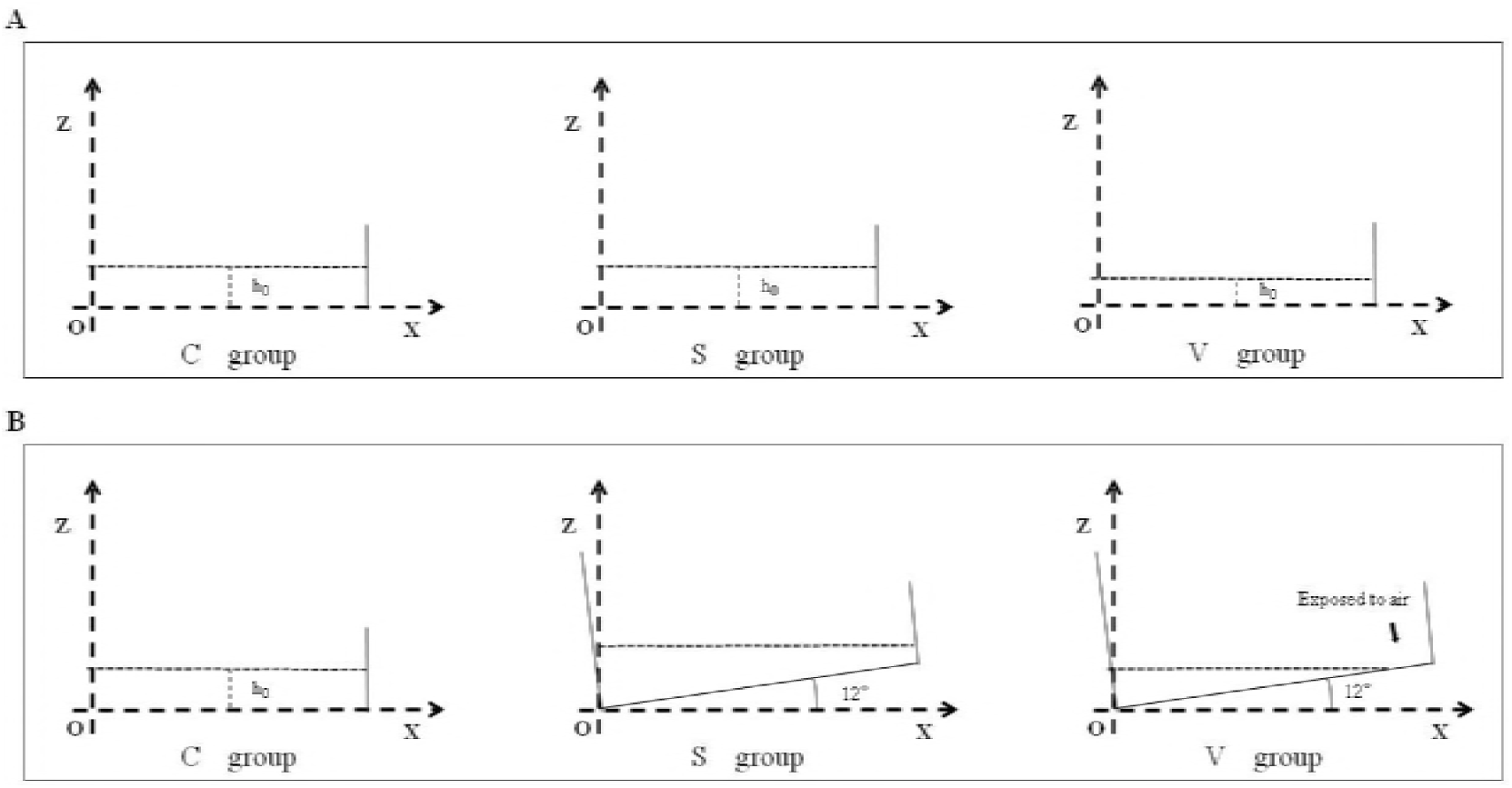
Set up a middle vertical cross-section at culture well. (**A**) The conditions of three groups when the table concentrator was in horizontal position. (**B**)The different conditions occur in the three groups when the table concentrator was 12° inclined.

In the C group, each well was filled with 3.5ml medium and the h_0_ was 3.6mm. The culture plate were settled on horizon plane with no stimulation.

In the S group, each well was filled with 3.5ml medium and the h_0_ was 3.6mm. When the table concentrator moved from horizon plane to 12°, the entire dish bottom always covered by medium. As a result, no area would be exposed during the interference process. The FSS would not be cut, and the variation will be constantly and regularly. FSS changing was more stable compared with the V group. So, the FSS in S group was defined as stable changing.

In the V group, each well was filled 2.5ml medium and h_0_ was 2.604mm. When the table concentrator moved from horizon plane to 12°, an area of about 95.28mm^2^ at the peripheral regions were exposed to air (Figure 2). Because the oscillating was cyclical and symmetrical.,there were two 10% peripheral regions of well bottom exposed. The FSS was dramatically changed at this exposing moment. So, the FSS in the V group was defined as unstable changing.

**Figure 2.**
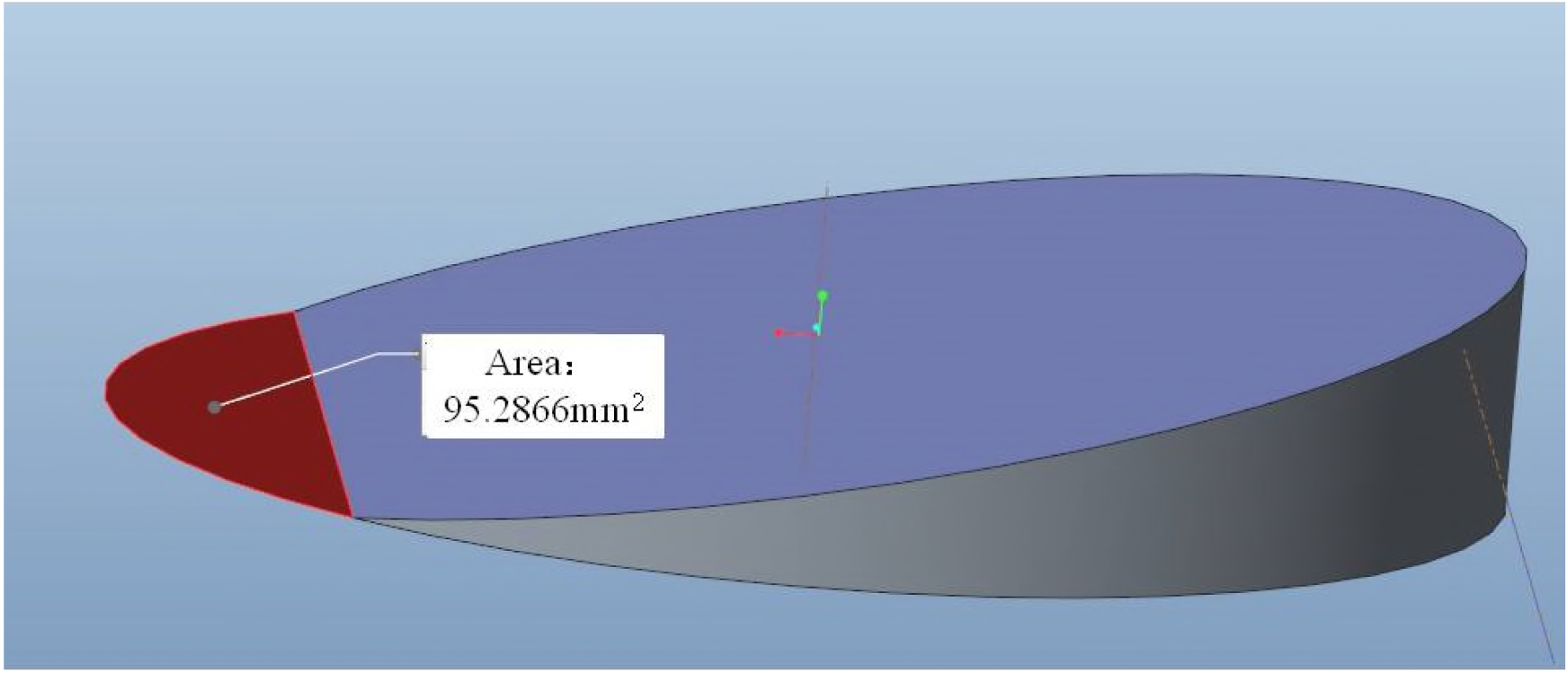
In the V group, an area about 95.282mm^2^ in peripheral regions was exposed to air that was calculated using Pro-Engineer 5.0 software.

As mentioned above, the FSS was an non uniform pattern where the stress were changing all the time. It was too difficult to estimate the FSS in every point and every moment. As a result, the shear stress at the center of dish bottom when the dish was horizontal during the oscillation process was defined as characteristic 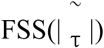. Because the FSS variation would around this value, it could be seemed as a standard to estimate FSS in S group and V group. According to the equation 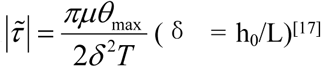 ^[17]^, the characteristic 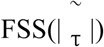 in the S group was 0.62dyn/cm^2^ and in the V group was 1.18dyn/cm^2^. The variation of FSS at three point(x/L=0.25,0.5,0.75) in the two groups of different time was in Figure 2. The peak FSS at different point in the two groups was in Figure 3.

**Figure 3.**
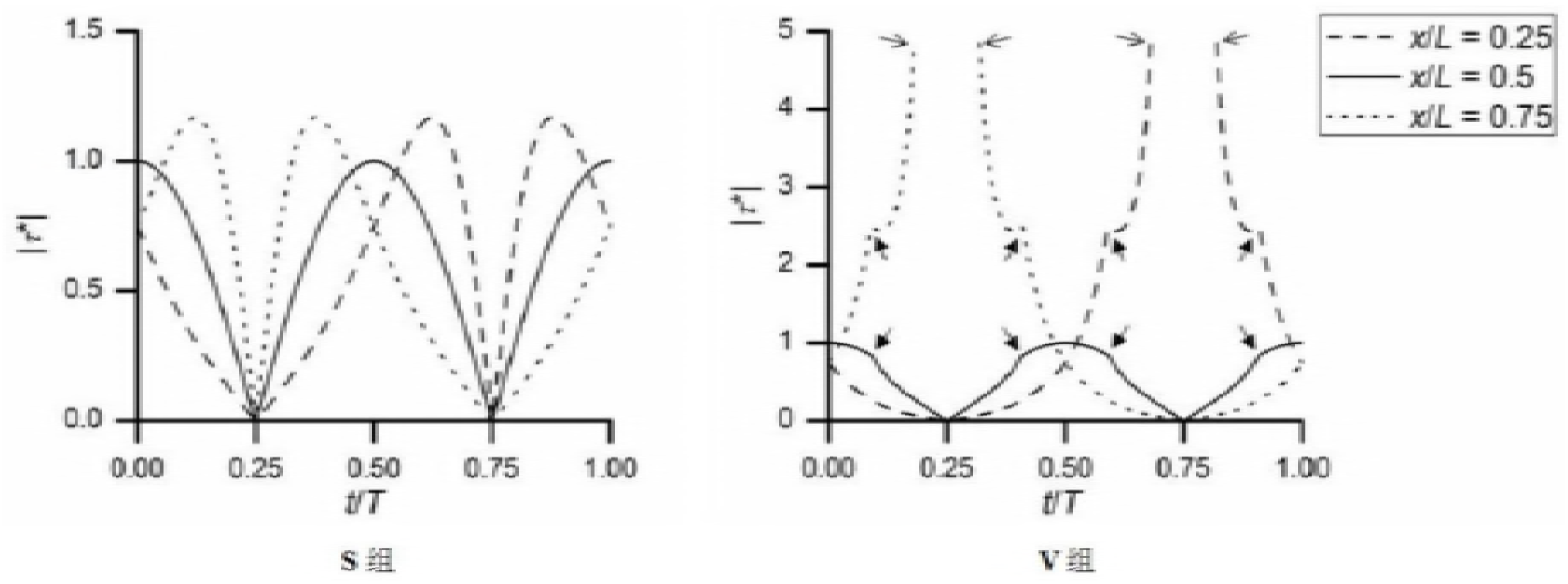
The variation of FSS at the three point(x/L=0.25,0.5,0.75) in the two groups^[17]^.(|τ^*^ means the ratio of FSS to characteristic 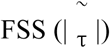

### 3.2 Expression of OPG, RANKL, ALP, OCN, RUNX2 and RANK mRNA

PCR results showed that osteogenesis in S group was prominent than other groups. Some mRNA which was associated with osteogenesis were detected on the 3^rd^ day. RANKL mRNA expression in the S group was in a lower level compared with the C group. The OPG mRNA expression in the S group was significantly greater than the C group. Only the S group had a significant difference with the C group in RANKL/OPG mRNA expression. The change of RANKL and OPG mRNA expression in the V group was not significant. RANK was associated with osteoclastgenesis. The mRNA expression in the S group was much lower versus the C group, but its expression in the V group was not significantly different with other groups. The mRNA expression of RUNX2, OCN, ALP were shown in the diagram. Only the RUNX2 had significant differences between the S group and the C group.

### 3.3 Mineralized nodules

Mineralized nodules represent the osteogenesis level of osteoblasts. We quantified mineralized nodules for number where the area was ≥100µm^2^, ≥500µm^2^ and ≥1000µm^2^. As well as quantified for total areas respectively. Both in the areas ≥1000µm^2^ and ≥500µm^2^, the number in the S group were much more than the other groups. In the case of ≥100µm^2^, the S group was much more than the C group, and no significant change compared with the V group. Similarly, the total area of the mineralized nodules in the S group was largest amongst the three groups.

### 3.4 TRAP Staining

Osteoclast differentiation was evaluated by the marker tartrate-resistant acid phosphatese(TRAP). TRAP is the main absorptive function protein of BMCs. The area of TRAP+ cell represents the differentiation level of osteoclasts. Comparing the TRAP+ area among the three groups, we found the areas was smallest in the S group. It indicated that stable changing FSS can inhibited osteoclastsgenesis.

## 4. Dicussion

Bone can be defined as a homeostasis tissue that consists of osteoclast and osteoblast dynamic equilibrium. In the continuous cycle, the osteoclast is responsible for absorbing, dissolving both the inorganic and organic components of the matrix by secreting protons and collagenolytic enzymes ^[18]^. Osteoblast is responsible for bone remodeling and formation ^[20]^. Furthermore, osteoclasts interplay with osteoblasts at different differentiation stages ^[21]^.

Many studies have been focused on either promoting osteoblast activity or inhibiting bone resorption to avoid bone loss, through hormones or chemical compounds to disrupt specific parts of the remodeling cycle^[22]^. However these studies overlooked the intrinsic ability of bone tissue to adapt to external stimuli from the environment. Actually, mechanical loading can help to enhance bone mass by promoting bone formation and suppressing bone resorption in vivo^[23]^.Physiologically, shear stress can act as a mainly mechanical stimuli from blood flow and interstitial fluid to cells ^[24]^. Many studies have presented that adaptable FSS can improve osteogenesis, like 0.15dyn/cm^2[25]^, 12 dyn/cm^2 [26–28]^, 16dyn/cm^2[29]^, and 12 × 10^-5^N^[30]^. Zhao, LG proved that fluid shear stress promotes ALP activity and the expression of OPN and OCN through MEK5/ERK5 pathway^[31]^. Besides that, primary murine osteoblasts were subjected to FSS in 5 Hz and 10 dyn/cm^2^, RANKL was induced by 2 hours post-FSS and keep elevating for at least 8 hours^[32]^. Although there are various studies have fond the relationship between FSS and bone cells, most methods supplied a sustainable FSS to osteoblast or BMCs to study the effects in different FSS magnitude. They created a model to supply intermittent FSS to bone cells to explore the relationship between FSS and time. However, all above FSS is just keeping at a certain values, that cannot imitated the FSS in real conditions around BMCs in vivo. Actually, FSS is not constant in vivo based on different physical activities. In vivo testing of the human femur, the pressure reached 1 to 2 kPa during physiologic loading with, the higher pressure in head than neck, and it goes up with load rate^[33]^. Besides, the effect was decreasing over time. Cui L et. al confirmed a constantly FSS can enhance OPG mRNA expression at 12 hours, but no significant difference at 24 hours^[11]^, indicate that cells can gradually adapt to mechanical intervention when the stimulus is constant^[34]^.

Although osteocytes are the mainly mechanosensory cells in bone tissue, but those cells in the bone marrow are also mechanically sensitive. C.P.Soves conducted a mechanical loading on mice limb in vivo and that induced the mechanical signal between bone marrow cells^[7]^. Li and Mantila Roosa proved that mechanical loading can increase proliferation and osteogenic gene expression of marrow stromal cells^[35, 36]^. Besides, osteoblast and osteoclast interplay with each other to keep dynamic equilibrium of bone tissue at bone marrow environment. There are at least three approachs in osteoclast-osteoblast communication. First, osteoblasts and osteoclasts can have a direct contact, allowing membrane-bound ligands and receptors to interact. Second, they form a gap junctions allowing passage of small water-soluble molecules between the two cell types. Third, communication occurs to diffuse paracrine factors, such as growth factors, cytokines, chemokines and other small molecules ^[37]^. The cell-cell communication occurred at basic multicellular unit (BMU) in bone marrow, a group of cells that achieve structural changes and skeletal renewal by the process of bone remodeling^[38]^. The communication occurred at every phases of bone remolding. At initiation phase of remolding process, hematopoietic precursors are recruited to the BMU and then differentiate into osteoclasts^[39]^. In this phase, OPG-RANKL-RANK axis and LGR4-RANKL-RANK axis plays an important role in regulation of osteoclastgenesis. RANKL and RANK are indispensable factor in osteoclast differentiation. OPG and LGR4 will compete with RANK to bind with RANKL. It directly influences the differentiation of hematopoietic precursors to osteoclast^[37, 40, 41]^. OPG is produced by many types of cells like osteoblast and osteocyte. In bone marrow, it has been shown that B cells are the major source of OPG in mice^[42]^. Besides that, M-CSF is also the key factor in osteoclast differentiation. Osteoblast can secrete M-CSF and MCP1 to keep osteoclast precursors proliferation and surviv in bone marrow. In addtion, Sema3A/Nrp^[43]^ and Lysophosphatidic acid (LPA)^[44]^ are released by osteoblast can affect osteoclast differentiation in different pathway. Subsequently, transition from bone resorption to formation by osteoblast can also be mediated by osteoclast. It directly influences the differentiation and activation of osteoblasts in resorbed lacunae to refill it with new bone. Most of these molecules are released by resorbed bone matrix or osteoclast. Matsuoka founds that Complement Component 3a (C3a) was progressively increased during osteoclast differentiation. Its bioactive fragment 3a can stimulate the osteoblast differentiation. The C3a receptor was mainly expressed in calvarial osteoblasts^[45]^. Negishi-Koga showed that osteoclasts can highly express Sema4D and it potently inhibits bone formation through the receptor plexin-β1 of osteoblasts^[46]^. Besides, some factors also serve as a bridge for the communication. Estrogen can affect osteoclast survival through the upregulation of FasL in osteoblasts, inducing the apoptosis of osteoclasts precursors^[47]^. Considering the interaction between osteoclast and osteoblast in bone marrow, we thought that choosing BMCs was appropriate for studying the specific effect of FSS in bone homeostasis.

As the table concentrator tray incline from horizonal plane to maxium angle, the medium depth varied as time goes, that will led to a continuous change of during the process. It gave cells a variable mechanical stress as an intervention. Considering the different changing ways of FSS, we set two different cases in FSS. In the first case (S group), there were enough medium in the well. The entire dish bottom could be covered by medium in whole oscillating process. In the other case (V group), there were not enough medium in the well. With the angle of culture plate reached 12°, medium would flow to one side. Some peripheral area in the other side would be exposed to air. When the area exposed, the FSS dramatically changed. Xiaozhou Zhou proved that if the oscillating angle is adaptable or the medium in the plate is enough, the FSS variation will be regularly^[17]^. Considering 3ml is a common volume of medium for standard six-well culture plate. In order to ensuring sufficient nutrients for cells growth, we set 3.5ml medium as the S group and 2.5ml medium as the V group to satisfy our research demands.

After the 3^rd^ day of intervention, we found that the osteogenesis in S group was best of all the groups, and osteoclastgenesis was worst in the three groups. We found that the RUNX2, OPG, and RANKL/OPG in S group have significant difference with C group in our studie. RANK and RANKL are mainly factors of osteocalstgensis, their combination can promote the differentiation of osteoclast. In OPG-RANKL-RANK pathway, OPG combines with RANKL then block the combination of RANK with RANKL ^[49]^,so OPG also name as a osteoclastogenesis inhibitory factor. The increasing of OPG can lead decreasing of osteoclastgensis directly. RUNX2 is a regulator in osteoblast differentiation, it plays a role in bone formation. All of the results indicated that stable changing of FSS improved the osteogenesis in BMCs and that can be detected on the 3^rd^ day during FSS stimulation. All the results of osteoblast associated with mRNA expression, only ALP and OCN have no significantly difference. ALP is a bone formation markers and OCN is a factor of bone metabolism and energy metabolism, both of them are usually being found at later stage of osteoblast differentiation and functioning in osteogenesis. In our studie, FSS magnitude was relatively lower, and three days was too short to detect the variation of ALP and OCN mRNA levels. On the 6^th^ day, we use minerlization nodules as a sign to detect the degree of osteogenesis. As we predicted, the result in S group was remarkable different from C group. The above results demonstrated that stable changing FSS has a positive effect on osteoblast formation.

Beyond that, we also investigate the degree of osteoclastgenesis through detection of RANK mRNA expression level and TRAP+ stain. RANK is the major regulator of ostelclastgensis and the down steam of RANKL^[39]^. RANKL contact with RANK to recruit TNF receptor associated factors (TRAFs) to activate various signaling pathways to promote osteoclastogenesis. But their contact can be impaired by OPG occupying the bind site. The decreasing of RANK mRNA expression means the osteoclastgenesis are inhibited. At RANK mRNA expression level, the S group was lower than the C group. The total area of TRAP + stain in the S group was the smallest among three groups, it proved that stable changing FSS also can inhibit osteoclastgenesis through decrease RANK mRNA expression. Besides, most of these detection results of the V group has no difference with the control group. However, we still noticed the small change of V group. Ban Y proved that compared with intermittent flow profusion, the continuous flow is a better environment for osteoblast activity ^[54]^. That indicated the effect from unstable changing FSS was more weakly to stable changing FSS. In conclusion, we thought that enough volume of medium can afford a stable changing FSS, and this was an appropriate stimuli for osteogenesis in BMCs.

## 5. Conclusion

This FSS device perfectly mimicked the microenvironment in vivo and it has effect in osteogenesis at complex marrow cells culture system. Enough volume of culture medium produced a stable changing FSS, and it was beneficial to osteogenesis. Considering the effects of FSS on BMCs, it is meaningful to apply such FSS to osteoblast or osteoclast, and it can help to reveal the specific signal pathway from FSS to cells.

**Figure 4.**
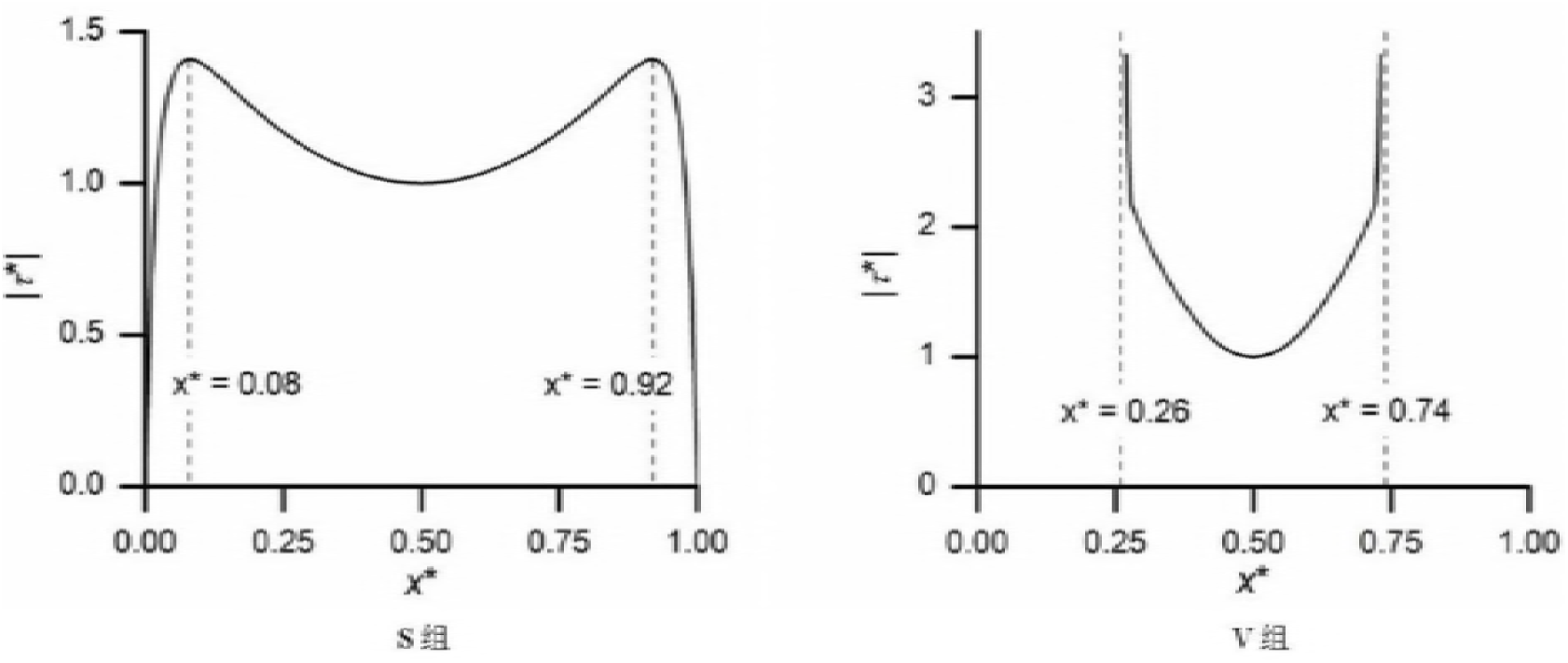
The peak FSS at different point in the two groups^[17]^.(x^*^=x/L; |τ^*^| means the ratio of FSS to characteristic 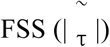

**Figure 5.**
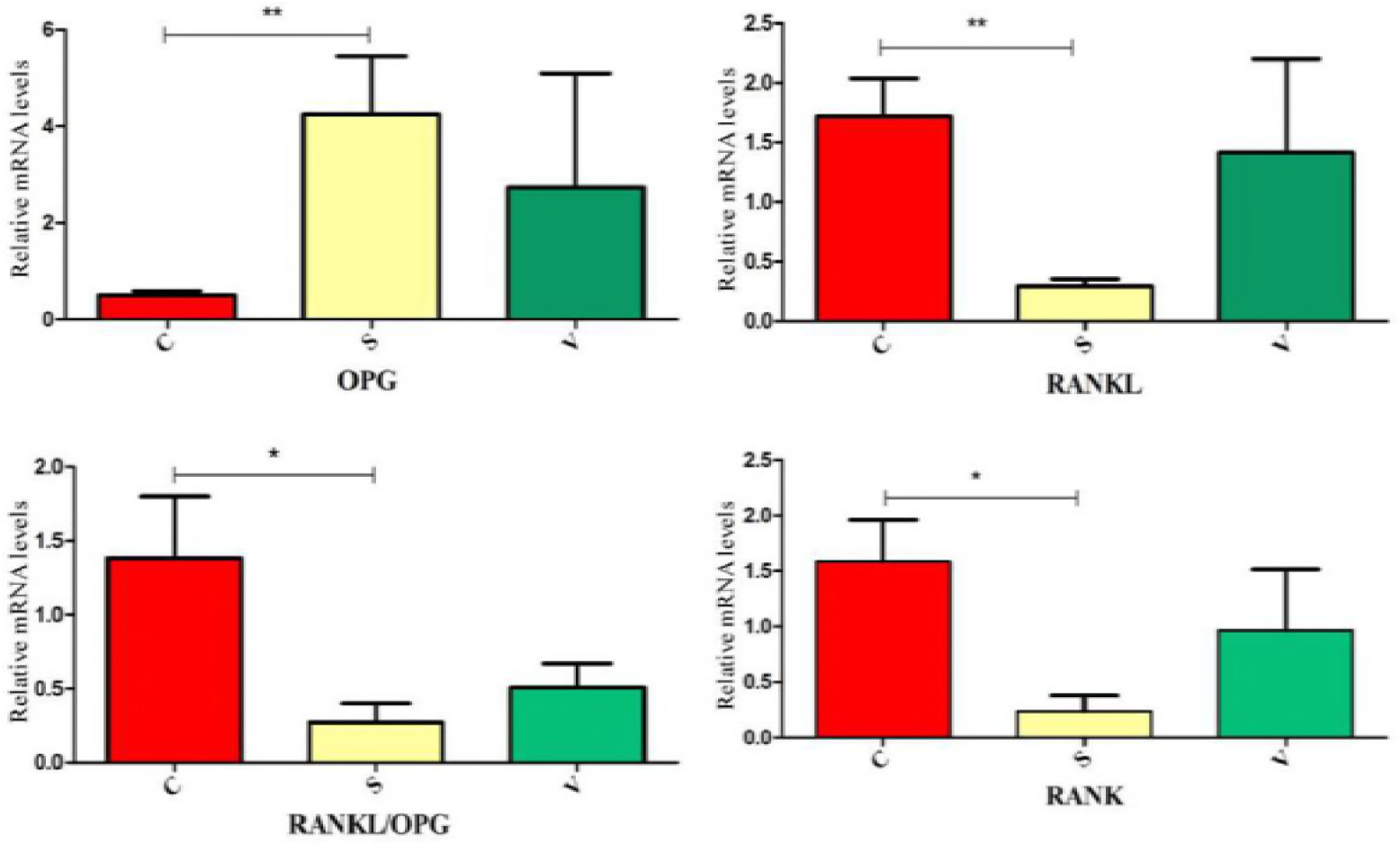
The relatively mRNA expression of RANKL, OPG, RANKL/OPG and RANK were showed in the diagram. \**p<0.05;* \*\**p<0.001*

**Figure 6.**
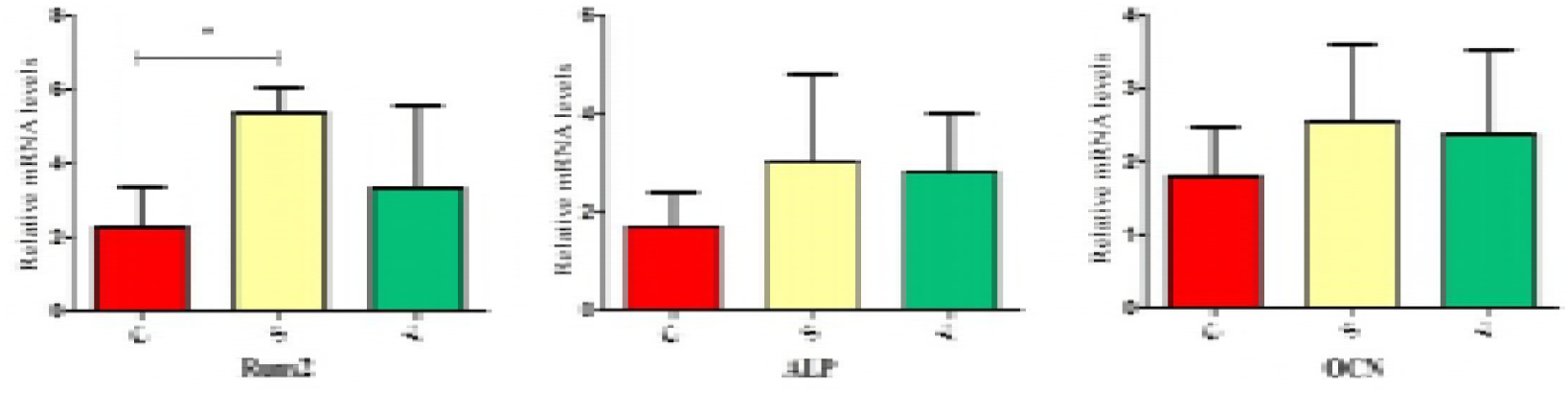
The relatively mRNA expression of RUNX2, OCN and ALP were showed in diagram. \**p<0.05*

**Figure 7.**
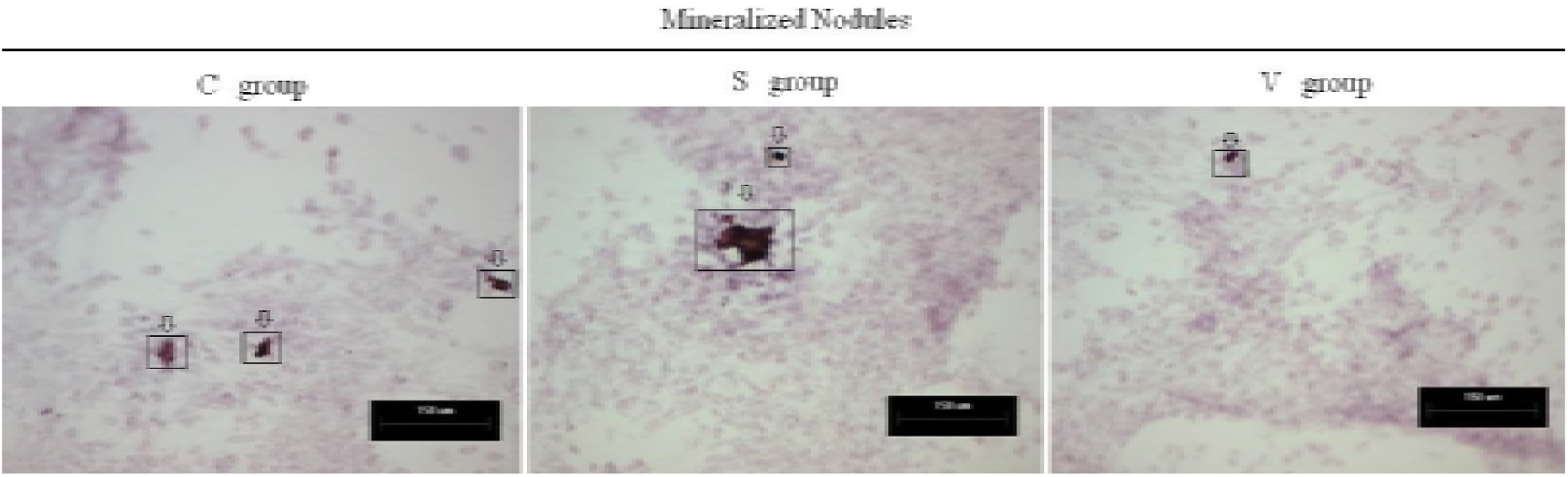
The formed mineralized nodules were stained positively with alizarin red S stain and were represented by the arrows.

**Figure 8.**
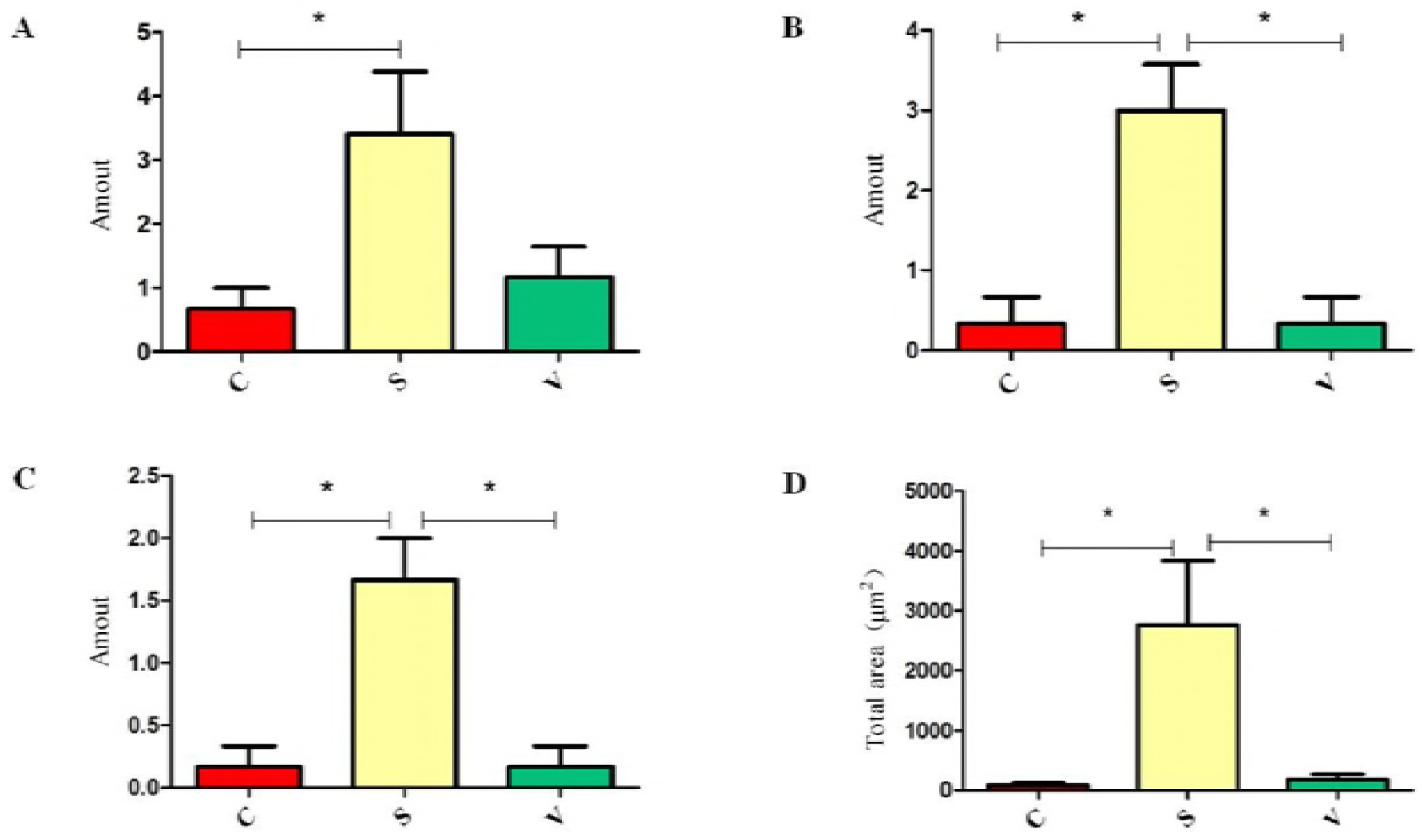
The amount of mineralized nodules in different areas in the three groups.(A) The amount of mineralized nodules area ≥100μm^2^.(B) The amount of mineralized nodules area ≥500μm^2^. (C) The amount of mineralized nodules area ≥1000μm^2^. (D) The total area of mineralized nodules.\**p<0.05*

**Figure 9.**
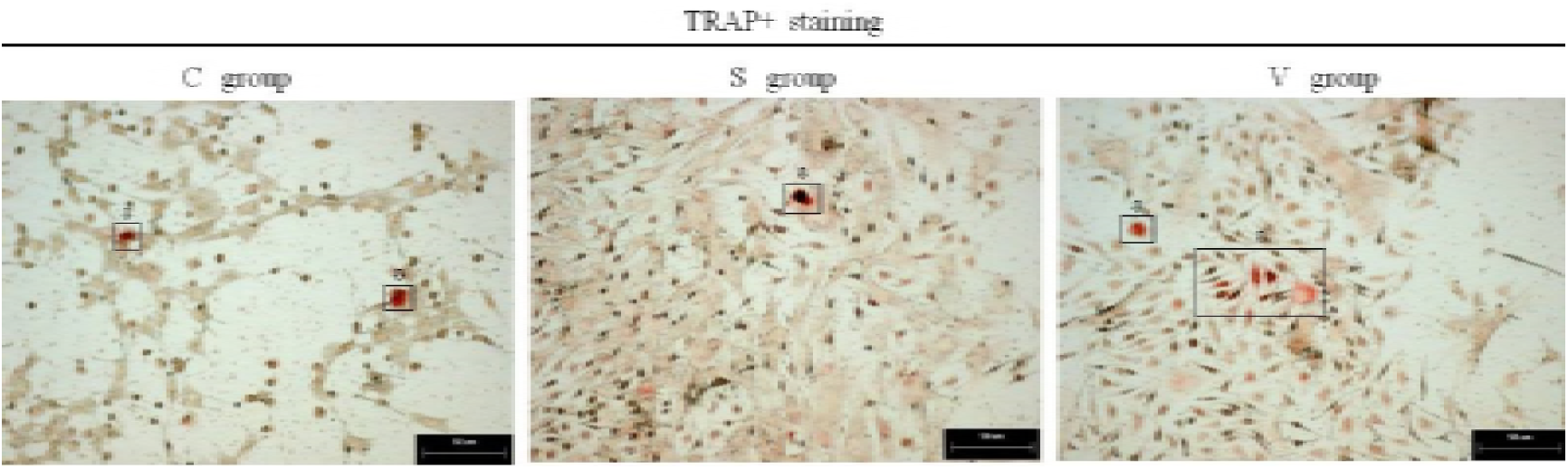
The osteoclast cells were stained positively with TRAP+ staining and were represented by the arrows.

**Figure 10.**
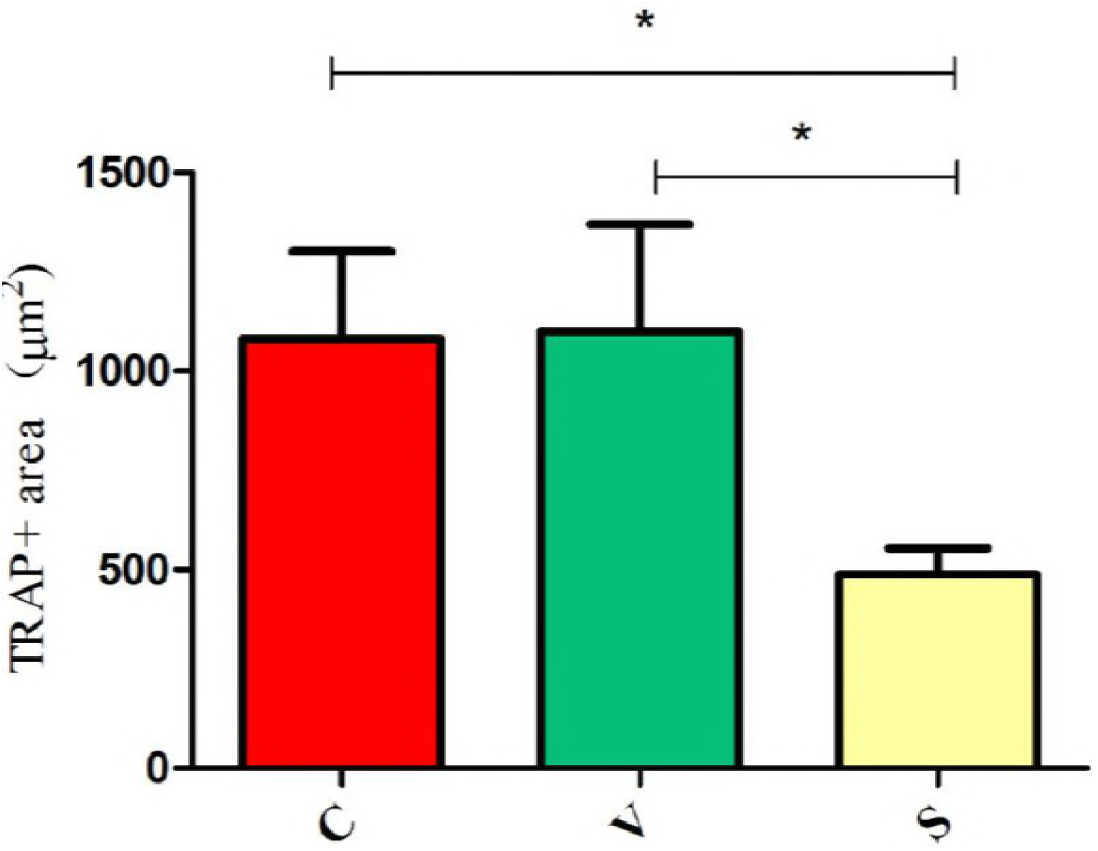
The total area of TRAP + cells in three groups were showed in diagram. \**p<0.05*

